# A genome-wide survey of DNA methylation in *Panax notoginseng* reveals CHH hyper-methylation regulates the after-ripening and dormancy of recalcitrant seeds

**DOI:** 10.1101/2023.12.05.570139

**Authors:** Na Ge, Jin-Shan Jia, Qing-Yan Wang, Chao-Lin Li, Min Huang, Jun-Wen Chen

## Abstract

DNA methylation plays a crucial role in regulating fruit ripening and seed development. It remains unknown about the dynamic characteristics of DNA methylation and its regulation mechanisms in morpho-physiological dormancy (MPD)-typed seeds with recalcitrant characteristics. The *P. notoginseng* seeds are defined by the MPD and are characterized by a strong sensitivity to dehydration during the after-ripening process. We performed DNA methylomes, siRNA profiles, and transcriptomes of embryo and endosperm in *P. notoginseng* seeds at different after-ripening stages. Herein, we find that the CHH hyper-methylation contributes to the global increase in DNA methylation during the after-ripening process of *P. notoginseng* seeds. The endosperm genome is hyper-methylated compared to the embryo genome. The CHH hyper-methylation is caused by the high expression level of DNA methyltransferase *PnCMT2* in the embryo, and *PnDRM2* in the endosperm, respectively. The CHH hyper-methylation alters gene transcription levels to regulate the after-ripening and dormancy of recalcitrant seeds. For example, it inhibits the expression of genes in embryo development to make seeds maintain a dormant status, whereas it activates the expression of genes in the hormone-mediated signaling pathway, and energy metabolism to accomplish the MPD-typed seed after-ripening process. Together, our findings reveal a global increase in DNA methylation and its vital driver in gene expression, and thus elucidate how global CHH hyper-methylation regulates the after-ripening in recalcitrant MPD-typed seeds. This work establishes a key role for epigenetics in regulating the dormancy of MPD-typed seeds with recalcitrant characteristics.

## Introduction

Cytosine DNA methylation is an important epigenetic modification in plants(Zhang *et al*., 2018). It is involved in regulating genetic functions, including transcription, replication, DNA repair, and cell differentiation, thus playing an essential regulatory role in plant growth, development, and evolution(Slotkin *et al*., 2009). DNA methylation exists in the symmetric CG and CHG sequence contexts and in the asymmetric CHH sequence contexts (H stands for A, T, or C)(Zhang *et al*., 2006; Deleris *et al*., 2016). The mechanisms of DNA methylation consist of maintenance methylation and de novo methylation(Law & Jacobsen, 2010). CG and CHG methylation is supported by DNA methyltransferases METHYLTRANSFERASE 1 (MET1) and CHROMOMETHYLASE 3 (CMT3), respectively, and CHH methylation is done by CMT2(Deleris *et al*., 2016). De novo DNA methylation cloud be established by DOMAIN REARRANGED METHYLTRANSFERASES (DRM1 and DRM2) through the RNA-directed DNA methylation (RdDM) pathway(Wang *et al*., 2022). In the RdDM pathway, DNA-dependent RNA polymerases Pol IV generates short P4-RNAs (26 to 50 nt) that are converted into double-stranded RNA (dsRNA) by RDR2, and are subsequently processed into 24-nt siRNAs by DICER-LIKE proteins 3(DCL3)(Zhai *et al*., 2015; Yang *et al*., 2016; Ye *et al*., 2016). The 24-nt siRNAs are loaded into AGO4 or AGO6 and paired with complementary scaffold RNAs and nascent transcripts produced by Pol V(Xu *et al*., 2013). The resulting complex recruits the DNA methyltransferase DRM2 to catalyze CHH methylation(Ye *et al*., 2016). In *Arabidopsis*, DNA methylation levels are dynamically regulated by DNA demethylases REPRESSOR OF SILENCING 1(ROS1), DEMETER (DME), DEMETER-LIKE 2 (DML2), and DML3(Bouyer *et al*., 2017). Dynamic DNA methylation is critical for regulating gene expression and maintaining genome stability.

In plants, DNA methylation has been shown to regulate fruit ripening and seed viability(Zhang *et al*., 2018; Michalak *et al*., 2022). An overall reduction in DNA methylation levels is observed in tomato (*Solanum lycopersicum*) fruit ripening(Teyssier *et al*., 2008; Zhong *et al*., 2013; Liu *et al*., 2015). The increased expression of DNA demethylase DME-LIKE 2 (DML2) has been linked to the loss of 5-methylcytosine (mC) DNA methylation in a naturally COLOURLESS NON-RIPENING tomato(Manning *et al*., 2006). The methylation level of the promoter region of the *MdMYB10* gene in apple (*Malus×domestica*) anthocyanin-deficient yellow-skin somatic mutant is significantly higher than those in red normal apples, and the methylation level progressively increases alongside fruit development and ripening(El-Sharkawy *et al*., 2015). In non-climacteric strawberry (*Fragaria vesca*) fruit, siRNA involved in RNA-directed DNA methylation is reduced, thus resulting in DNA hypo-methylation during fruit ripening(Cheng *et al*., 2018). In contrast, global DNA hyper-methylation is found during the process of orange fruit ripening, which is caused by the decreased expression of *CsDMEs* and *CsDMLs.* Furthermore, the ripening-induced DNA hyper-methylation represses the expression of photosynthesis-related genes and activates the expression of genes involved in abscisic acid responses during the process of orange fruit ripening(Huang *et al*., 2019). The seeds have been classified as orthodox, intermediate and recalcitrant types based on their storage behavior(Angelovici *et al*., 2010). A research on orthodox *Acer platanoides* L. seeds(Plitta *et al*., 2014) has revealed that DNA methylation is significantly enhanced when the seeds are desiccated from a water content of 1.04 g·g^-1^ to 0.17-0.19 g·g^-1^. Similarly, with the increased DNA methylation, the viability of orthodox *Pyrus communis* L. seeds remains high and constant(Michalak *et al*., 2013). In contrast, an analysis of recalcitrant *Quercus robur* seeds has demonstrated that desiccation-induced osmotic stress results in a negligible increase in the level of DNA methylation(Plitta *et al*., 2014). These observations have suggested that DNA methylation has a broad role in fruit ripening and seed viability. However, a genome-wide survey of DNA methylation has not been investigated in the recalcitrant seed with an after-ripening process. Thus, the dynamic characteristics of DNA methylation and its regulation mechanisms are unknown in recalcitrant seeds during the after-ripening process.

Seed dormancy, germination, and longevity determine the start of plant life cycle(Nee *et al*., 2017), (Penfield, 2017; Soppe & Bentsink, 2020). Seed dormancy and germination is a complex developmental process that is under strict genetic, epigenetic, and hormonal controls(Sun *et al*., 2020; Tognacca & Botto, 2021; Tuan *et al*., 2021). Different types of dormancy occur in seeds, and they are defined as physiological dormancy (PD), morphological dormancy (MD), and morpho-physiological dormancy (MPD) types. After-ripening is a process in which MPD-typed seeds experience a process of gradual reduction in dormancy level(Finch Savage & Leubner Metzger, 2006; Baskin & Baskin, 2007). DNA methylation regulates seed dormancy and after-ripening development(Nakabayashi *et al*., 2005; Qin *et al*., 2010). In angiosperms plants, the embryo and endosperm are produced by the double fertilization of egg cells and central cells, respectively, and the development of embryos and endosperm is accompanied by DNA methylation reconfiguration(Jullien *et al*., 2012). Endosperm tissue is extensively demethylated through the high expression of DNA glycosylases Demeter (DME) or the down-regulation of DNA methyltransferase-1 (MET1) activity in central cells at pre-fertilization in *Arabidopsis* seeds(Gehring *et al*., 2009; Bauer & Fischer, 2011). During the development of PD-typed *Arabidopsis* seeds, the increased methylation at the CHH site in the embryo is dependent on a sustained increase in chromatin methyltransferase 2 (CMT2) and RNA-directed DNA methylation (RdDM) activity, while the endosperm holds a low level of DNA methylation(Bouyer *et al*., 2017). Similarly, DNA methylation levels are substantially lower in the endosperm than in the embryo of PD-typed rice (*Oryza sativa* L.) seeds, and this is related to the activated expression of starch synthesis genes and glutenin precursor genes in the endosperm(Zemach *et al*., 2010). In *Arabidopsis* seeds, a global increase of CHH methylation is observed during seed development, while DNA methylation shows a drastic loss during seed germination(Bouyer *et al*., 2017). These dynamic changes in methylation level depend on RNA-directed DNA methylation pathways(Kawakatsu *et al*., 2017). The tissue-specific development of embryo and endosperm is regulated by the activity of DNA methylesterases and demethylases. However, it is unknown whether the dynamic DNA methylation observed in the model system also occurs in the MPD-typed seeds, and how DNA methylation changes in the embryo and endosperm during the recalcitrant seeds after-ripening.

*Panax notoginseng* (Burkill) F. H. Chen (Sanqi in Chinese) is a perennial herb in the family of Araliaceae(Chen *et al*., 2016), and it is an important and traditional medicinal plant, with a ∼2.3 Gb genome(Yang *et al*., 2021). Its seeds belong to the MPD group and have 45 ∼ 60 days of the after-ripening before germination. Moreover, *P. notoginseng* seeds show a typically recalcitrant characteristic that the seeds are highly sensitive to dehydration during the after-ripening process(Duan *et al*., 2014). Generally, the moisture content of seeds is about 67.3% at harvest, and seed viability can be only maintained for about 15 days under natural conditions(Cui *et al*., 1993; Liao *et al*., 2016). It has been shown that an incompletely developed embryo could be the main reason for the dormancy of *P. notoginseng* seeds, and the embryo at a heart-shaped period has to be further differentiated and developed during the after-ripening process(Yang *et al*., 2018). Exogenous GA_3_ application could change the ratio of GA and ABA, and thus greatly facilitate the development of the embryo in recalcitrant *P. notoginseng* seeds(Ge *et al*., 2023). Likewise, the DNA methylation inhibitor, 5-azacytidine, might promote *P. notoginseng* fruit ripening and seed germination, thus suggesting that DNA methylation indeed regulates seed development and after-ripening-related biological processes(Huang *et al*., 2022). However, the pattern of global DNA methylation and the importance of DNA methylation have not been comprehensively elucidated in the MPD-typed *P. notoginseng* seeds during the after-ripening process.

In the present study, we performed DNA methylomes, siRNA profiles, and transcriptomes of embryo and endosperm in *P. notoginseng* seeds at different after-ripening stages, and our purpose is to investigate the epigenetic regulation of recalcitrant seed after-ripening and dormancy. We observed a globally increased DNA methylation during the after-ripening process. The DNA methylation inhibitor caused hypo-methylation and early germination of seeds, indicating that DNA methylation is vital for regulating the after-ripening of recalcitrant seeds. We also discovered that DNA hyper-methylation during the seed after-ripening is linked to the increased expression of DNA methyltransferase genes. Tobacco rattle virus (TRV)-induced gene silencing of DNA methyltransferase component, *PnCMT2*, led to early germination of *P. notoginseng* seeds. Remarkably, hundreds of seed development-related genes showed DNA hyper-methylation. Our results would help to explain the epigenetic regulation of recalcitrant seeds during the after-ripening process. In summary, our study firstly reveals the characteristic of the genome-wide dynamics of DNA methylation during the after-ripening process of recalcitrant seeds, and proposes a certain mechanism for maintaining the global CHH hyper-methylation to regulate the after-ripening in recalcitrant MPD-typed seeds.

## Materials and methods

### Plant materials

The *P. notoginseng* plants (Fig. S1) were grown in the experimental farm of Wenshan Miao Xiang *P. notoginseng* Industrial Co., Ltd., China (Longitude 104°32′, latitude 23°53′). Mature seeds of *P. notoginseng* (Fig. 1) were harvested from the plants of 3-year-olds in November. After being artificially peeled, the seeds were disinfected with a 5% CuSO_4_ bactericidal solution. The seeds were placed in a ventilated net basket for 50 days in a sandy stratification chamber (15 ± 5℃). Based on the previous morphological descriptions(Ge *et al*., 2023), seeds at three different after-ripening processes (CK, seeds harvested, 0 days after-ripening (DAR); T1, seed morphology completely developed, 30 DAR; T2, seed dormancy released, 50 DAR) were collected and dissected (Fig. 1). Embryo (Em) and endosperm (En) tissue of *P. notoginseng* seeds were collected and frozen in liquid nitrogen and kept at -80 °C for DNA and RNA extraction. Three biological replicates were used for each after-ripening process.

**Figure 1.**
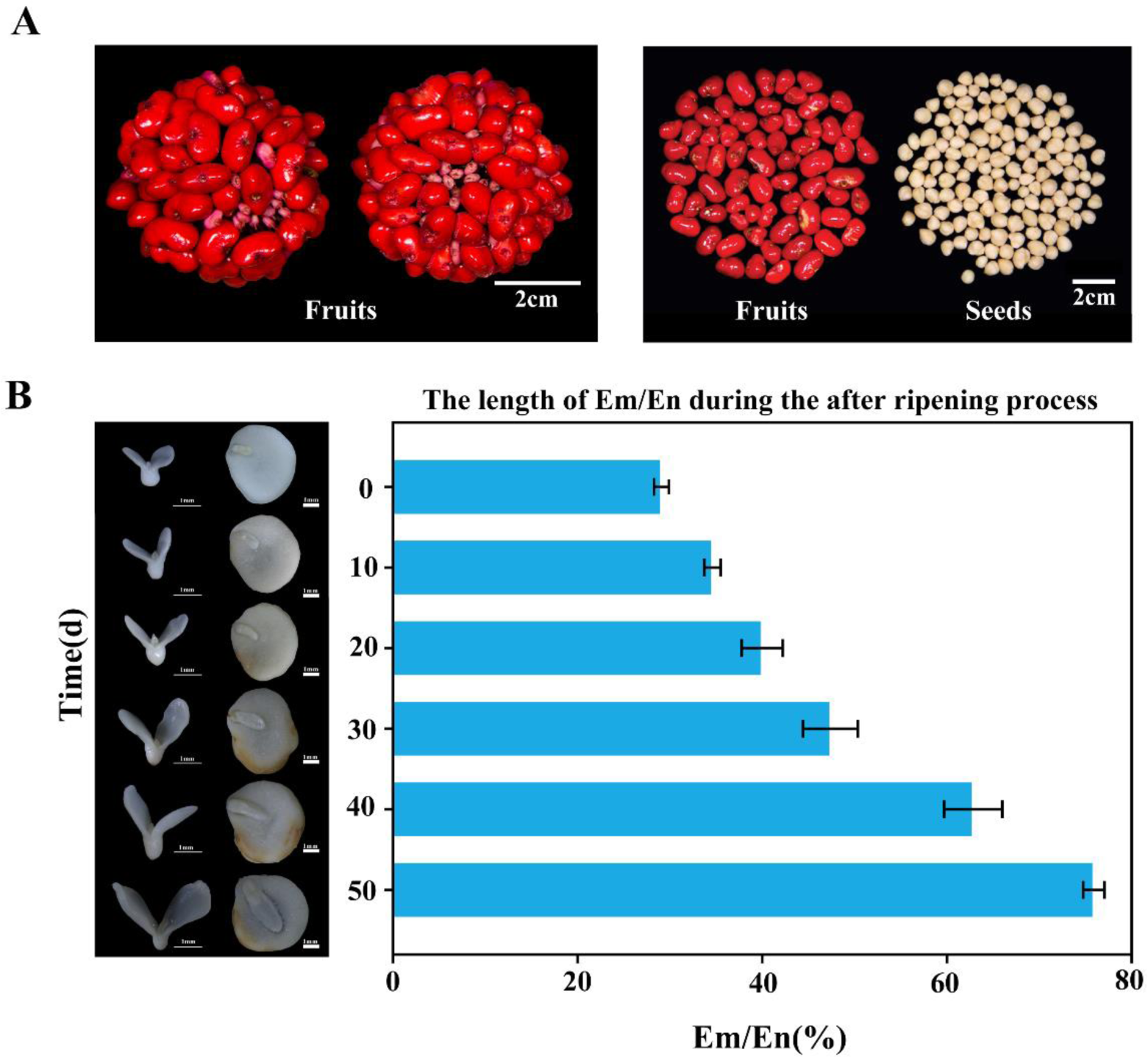
Stereoscopic micrographs of *P. notoginseng* fruit, seed sections (A), and the changes of the embryo (B), endosperm, and Em/En during the after-ripening process.

The seeds at 0 DAR were used for DNA methylation inhibitor and promoter treatments. In the treatment, the seeds were submitted to the concentrations 10 μmol L^-1^ by soaking for 24 h, 5-azacytidine and hydroxyurea (MedChemExpress) dissolved in ddH_2_0 with 0.01% Triton X-100 to10 μmol L^-1^, and ddH_2_O-soaked treatment was considered as the negative control (mock).

### Bisulfite sequencing and data analysis

The DNeasy Plant Maxi Kit (Tiangen) was used to extract genomic DNA from the embryo and endosperm. Briefly, genomic DNAs were fragmented into 100-300bp using Sonication (Covaris, Massachusetts, USA) and purification was performed using a MiniElute PCR Purification Kit (Qiagen). The genomic fragments were ligated to methylated sequencing adapters. The fragments with the adapters were converted to bisulfite using a Methylation Gold kit. (Zymo, USA). The converted DNA fragments were subjected to PCR amplification and sequenced on an Illumina HiSeq 2500 at Gene Denovo Biotechnology Co. (Guangzhou, China).

For the data analysis, the raw reads containing more than 10% of unknown nucleotides (N) and low-quality reads containing more than 40% of low-quality (quality score < 20) bases were first removed. The clean reads were mapped to *P. notoginseng* reference genome(Yang *et al*., 2021) using BSMAP-2.90(Xi & Li, 2009). Methylated cytosines were then retrieved and tested with the correction algorithm using a custom Perl script(Lister *et al*., 2009). The methylation level was calculated based on methylated cytosine percentage in the whole genome for each sequence context (CG, CHG, and CHH). Differential DNA methylation between the two samples at each locus was determined using methyl Kit-1.7.1(Akalin *et al*., 2012). Fisher’s exact test (*P* < 0.05) was used to define differentially methylated cytosines (DMCs). The DMRs were identified as previously described with minor modifications(Cheng *et al*., 2018). In brief, the minimum read coverage of methylation status for a base was set to four. A 200-bp sliding window was used to evaluate patterns of methylation in different regions of the genome. Windows with absolute methylation difference ≥ 0.25, 0.25, and 0.15 for CG, CHG, and CHH, respectively, and Benjamini-Hochberg corrected false discovery rates (FDRs) < 0.05 (Fisher’s exact test) were selected. To avoid a 200-bp window with few cytosines, the windows with at least five cytosines in each sample were selected. The significant regions in all three comparisons were conceded as candidates for further analysis. DNA methylation was visualized using IGV software(Thorvaldsdottir *et al*., 2013). DMR-associated genes were identified as genes with the closest DMR located within 2 kb upstream of the transcription start site (TSS) and 2 kb downstream of the transcription end site (TES).

### Transcriptome sequencing analysis

For the samples collected from the seed after-ripening process, RNA was isolated using a Trizol reagent kit (Invitrogen, Carlsbad, CA, USA) according to the manufacturer instructions. The RNA was sequenced on an Illumina HiSeq2500 by Gene Denovo Biotechnology Co. (Guangzhou, China). To get high-quality clean reads, low-quality sequences (Q < 20) and adapters were filtered by fastp-0.18.0(Chen, S *et al*., 2018). The filtered clean reads were mapped to the *P. notoginseng* genome(Yang *et al*., 2021) using HISAT2. 2.4(Kim *et al*., 2015). StringTie v1.3.1(Pertea *et al*., 2015; Pertea *et al*., 2016) was used to assemble the mapped reads from each sample in a reference-based approach. The abundance and variations of gene expression were used as an FPKM (fragment per kilobase of transcript per million mapped reads) value. Differential expression analysis was performed by DESeq2(Love *et al*., 2014) between two different groups (and by edgeR(Robinson *et al*., 2010) between two samples). The genes with the parameter of false discovery rate (FDR) < 0.05 and absolute fold change ≥ 2 were considered differentially expressed genes (DEGs). Functional enrichment analysis was performed using GO enrichment analysis and KEGG pathway enrichment analysis.

### Small-RNA sequencing and data analysis

Three biological replicates of embryo and endosperm in each after-ripening process of seed were used to prepare for small RNA analysis. Total RNA was isolated using a Trizol reagent kit (Invitrogen, Carlsbad, CA, USA). Polyacrylamide gel electrophoresis (PAGE) was used to enrich RNA molecules < 100nt. Then a standard small RNA library was prepared, and the libraries were sequenced using Illumina HiSeq Xten in Gene Denovo Biotechnology Co. (Guangzhou, China). Illumina adapters and low-quality bases (quality score < 20) or containing unknown nucleotides (N) were removed. The cleaned 24-nt reads were mapped to the *P. notogensing* genome and identified as siRNA clusters using BWA-0.7.17(Yang *et al*., 2021).

### Phylogenetic analysis

The DNA methyltransferase and DNA demethylase of *P. notoginseng* and *Arabidopsis* were performed by multiple alignments using the MATTF program(Katoh & Standley, 2013). Then the neighbor-joining method with 1000 bootstrap replicates in MEGA11(Tamura *et al*., 2021) was used to construct the phylogenetic tree.

### McrBC-qPCR analysis

Genomic DNA from mock, 5-azacytidine-treated, and hydroxyurea-treated samples were extracted using a Plant Genomic DNA Kit (TIANGEN, China). Then ∼200 ng of DNA was digested with McrBC (Takara, 1234A, Japan) for 18 h at 37°C, and qPCR was tested (Additional file 5: Table S10) using digested DNA as template. A parallel reaction was used as a negative control in which equal amounts of DNA were added to a buffer without the enzyme. Three biological replicates were used for each experiment.

### Virus-induced gene silencing

Gene function was examined using a virus-induced gene silencing (VIGS) system based on tobacco rattle virus (TRV)(Liu *et al*., 2002). According to the treatment protocol suggested by Birch-Smith et al.(2004), fragments of *PnCMT2* and *PnPDS* were PCR-amplified (Additional file 5: Table S10) from *P. notoginseng* cDNA and ligated with the pTRV_2_ vector. The generated plasmids were introduced into Agrobacterium tumefaciens strain GV3101. Agrobacterium culture was infiltrated into pTRV_1_ and pTRV_2_ or its derivatives were mixed at a 1:1 (v/v) ratio and culture of pTRV_1_+pTRV_2_-*PnCMT2* vacuum infiltrated into 0 DAR seeds of *P. notoginseng*, the infiltrated seeds were placed in a ventilated net basket for 50 days in a sandy stratification chamber (15 ± 5℃). The culture of pTRV_1_+pTRV_2_-*PnPDS* vacuum was infiltrated into 1-week-old seedlings of *P. notoginseng*. The infiltrated seedlings were incubated under dark conditions for 72 h and then grown in a culture room at 20°C, 18/6 h light/dark, 110 μmol μmol/m^2^·s. To indicate the spatio-temporal pattern of gene silencing, plants infiltrated with pTRV_1_+pTRV_2_-*PnPDS* were used as the positive control.

### Quantitative RT-PCR analysis

Using the same scheme described for RNA-Seq, total RNA was isolated from seeds of *P. notogensing* samples. First-strand cDNA synthesis and real-time PCR were performed according to the method suggested by Jia *et al*.(2023). The internal control was *GAPDH* (*GLYCERALDEHYDE-3-PHOSPHATE DEHYDROGENASE*)(Ge *et al*., 2023) and the 2^-ΔΔCT^ method(Livak & Schmittgen, 2001) was used to calculate relative expression values. Primers used in this analysis are listed in Table S9. Three biological replicates were performed for each gene, and at least three technical replicates were conducted.

### Statistical analysis

Three biological replicates were performed in the experiment. The SPSS 20.0 software (Chicago, IL, USA) was used for statistical analysis, SigmaPlot 10.0 and GraphPad Prism 8.0 were used for drawing, and variables are presented as mean ± SD (n=3). We calculated the least significant difference (LSD) and *P* < 0.05 was considered statistically significant.

## Results

### DNA methylome of recalcitrant *P. notoginseng* seeds

To investigate the DNA methylome of *P. notoginseng* seeds, the seeds at three different after-ripening stages: 0, 30, and 50 days after-ripening (DAR) (referred to as CK∼T2) were performed whole-genome bisulfite sequencing for both embryo and endosperm tissues (Fig. 1). The genome of *P. notoginseng* (∼2.3Gb) was used as a reference in our analyses(Yang *et al*., 2021). For each sample, approximately 90% of the reads have been mapped to the reference genome, with more than 85% coverage. The conversion rates were > 99.1% for each bisulfite-treated library (Table S1). The sequencing depths were sufficient for further analysis.

Globally, DNA methylation levels reached 31.79% in the embryo, which was lower than that in the endosperm (39.17%) (Fig. 2). In the embryo, the DNA methylation levels were 82.15%, 74.61% and 14.71% in the CG, CHG, and CHH sequence contexts, respectively (Fig. 3A). For the endosperm, we found that cytosine methylation levels were higher in the CG (90.31%) than those in the CHG (82.18%) and the CHH (22.09%) (Fig. 2). To characterize DNA methylation distribution across the genome of *P. notoginseng* seeds, the distribution of DNA methylation and gene/transposable element (TE) densities were examined. As shown in Fig. S2, DNA methylation was not uniformly distributed across the chromosomes. It was observed that the methylation density of the embryo and endosperm show the relative similarity of DNA methylation patterns around genes and TEs. Generally, the DNA methylation was mainly enriched in heterochromatin that contained fewer genes (Fig. S2). The genes were found to be preferentially located in the chromosome arms and were mainly dominated by CG and CHG methylation, whereas CHH methylation levels were similar to the density of the genes (Fig. S2, S3). In contrast, the TE was distributed more uniformly on chromosomes (Fig. S2, S3).

**Figure 2.**
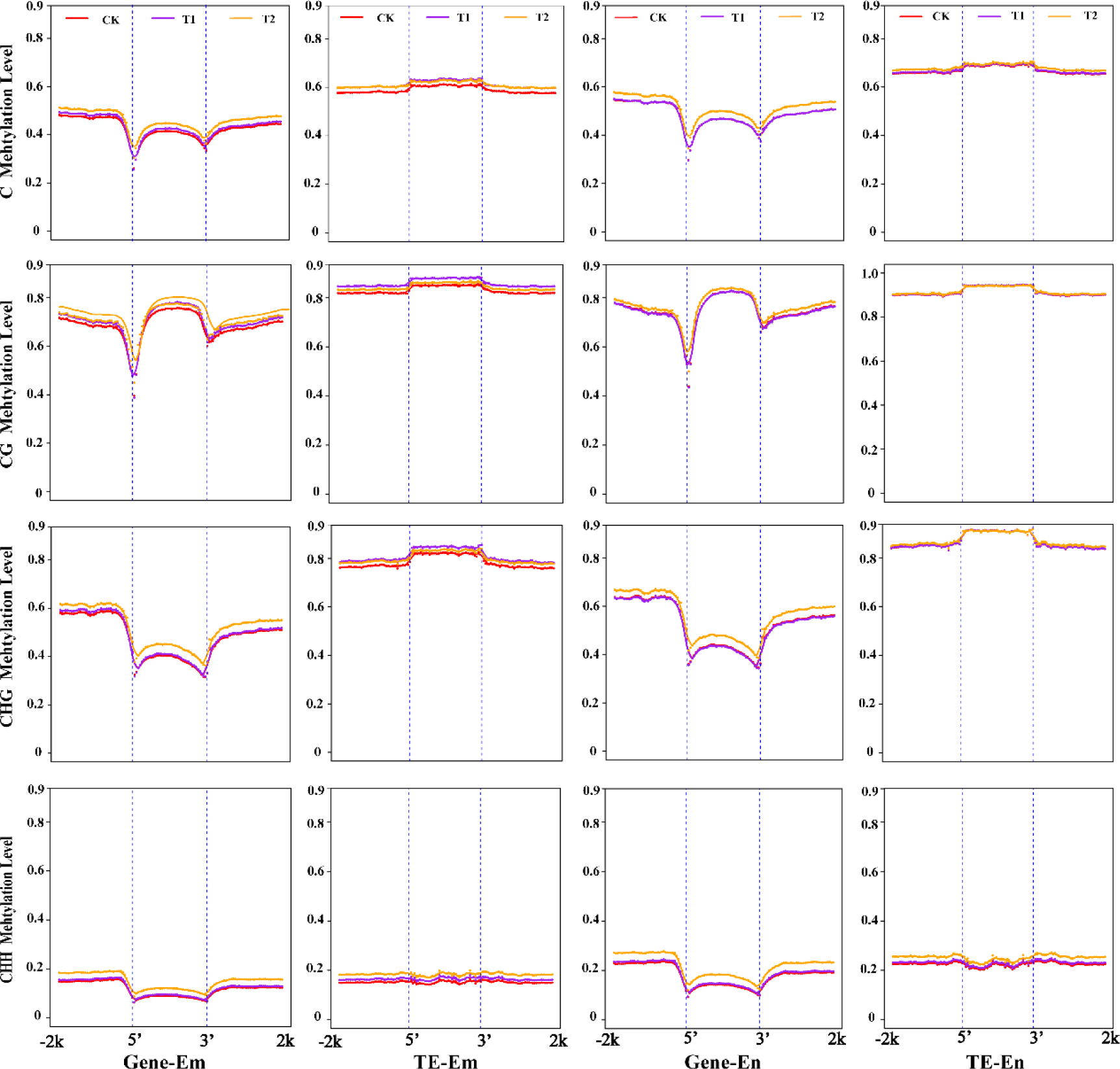
Characterization of the DNA methylations of embryo and endosperm during the after-ripening process of *P. notoginseng* seeds.

**Figure 3.**
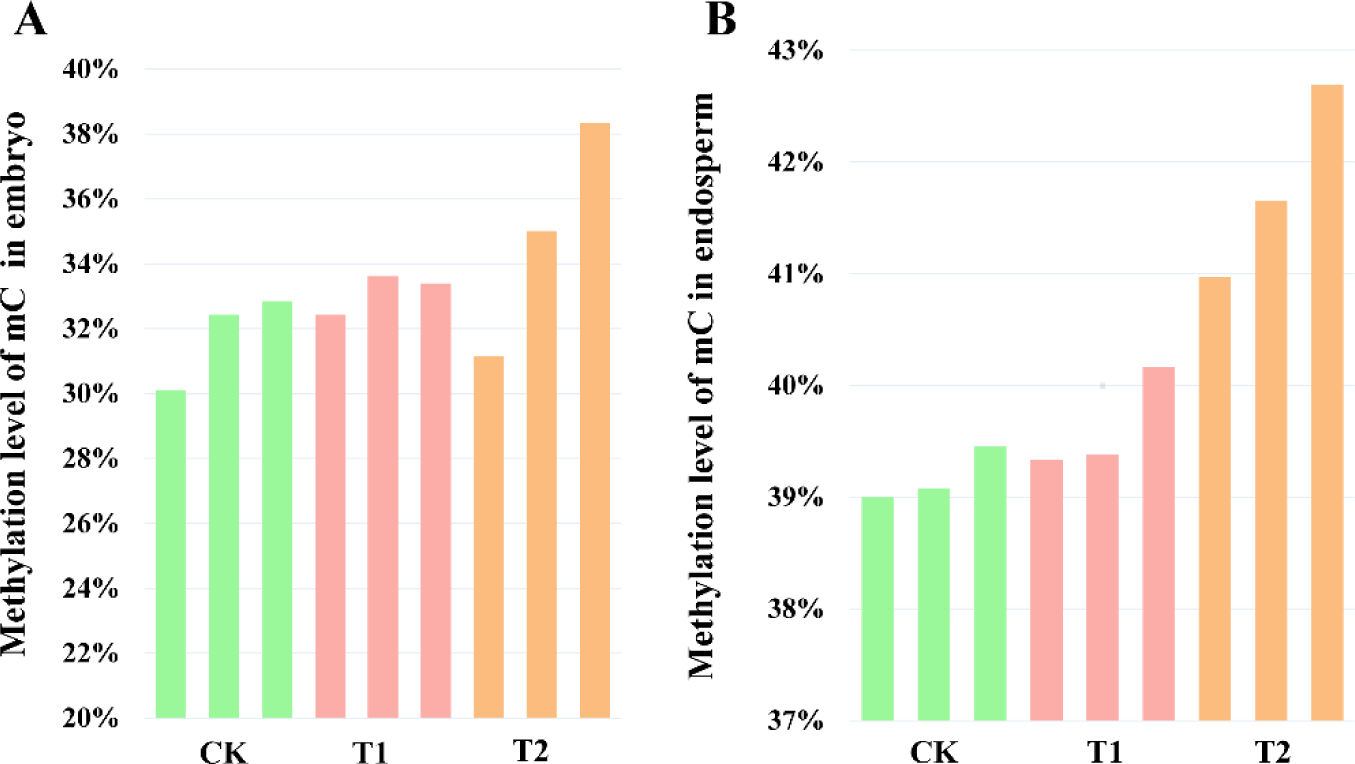
Genome-wide increase in DNA methylation in the embryo and endosperm during the after-ripening process (DAR). (A) DNA methylation in the embryo at 0, 30, and 50 DAR; (B) DNA methylation in the embryo at 0, 30, and 50 DAR.

### DNA methylation increases in the embryo and endosperm of *P. notoginseng* seeds during the after-ripening process

To investigate the dynamics of DNA methylation during the after-ripening process. We calculated the average DNA methylation levels at three different after-ripening stages (CK-T2) (Fig. S4, Table S4). It observed that the DNA methylation levels were enhanced from CK to T2 in the embryo (from 31.8 to 34.8%) and endosperm (from 39.2 to 41.2%) (Fig. 3), suggesting an increase in DNA methylation levels during the after-ripening process (Fig. S4). Additionally, the DNA methylation levels for genes and TEs were calculated in the embryo and endosperm during the after-ripening process. The gradual increases of DNA methylation are recorded in 5′ and 3′ flanking regions of genes and TE body and flanking regions (Fig. 2) from CK to T2. Among them, the most dynamic changes were observed in the body regions of the genes in the embryo and endosperm, whereas CG and CHG methylation in the TE of the endosperm maintained a high level during the after-ripening process (Fig. 2). Furthermore, the cause of the enhanced DNA methylation is largely attributed to the increase in CHH methylation (Fig. 4).

**Figure 4.**
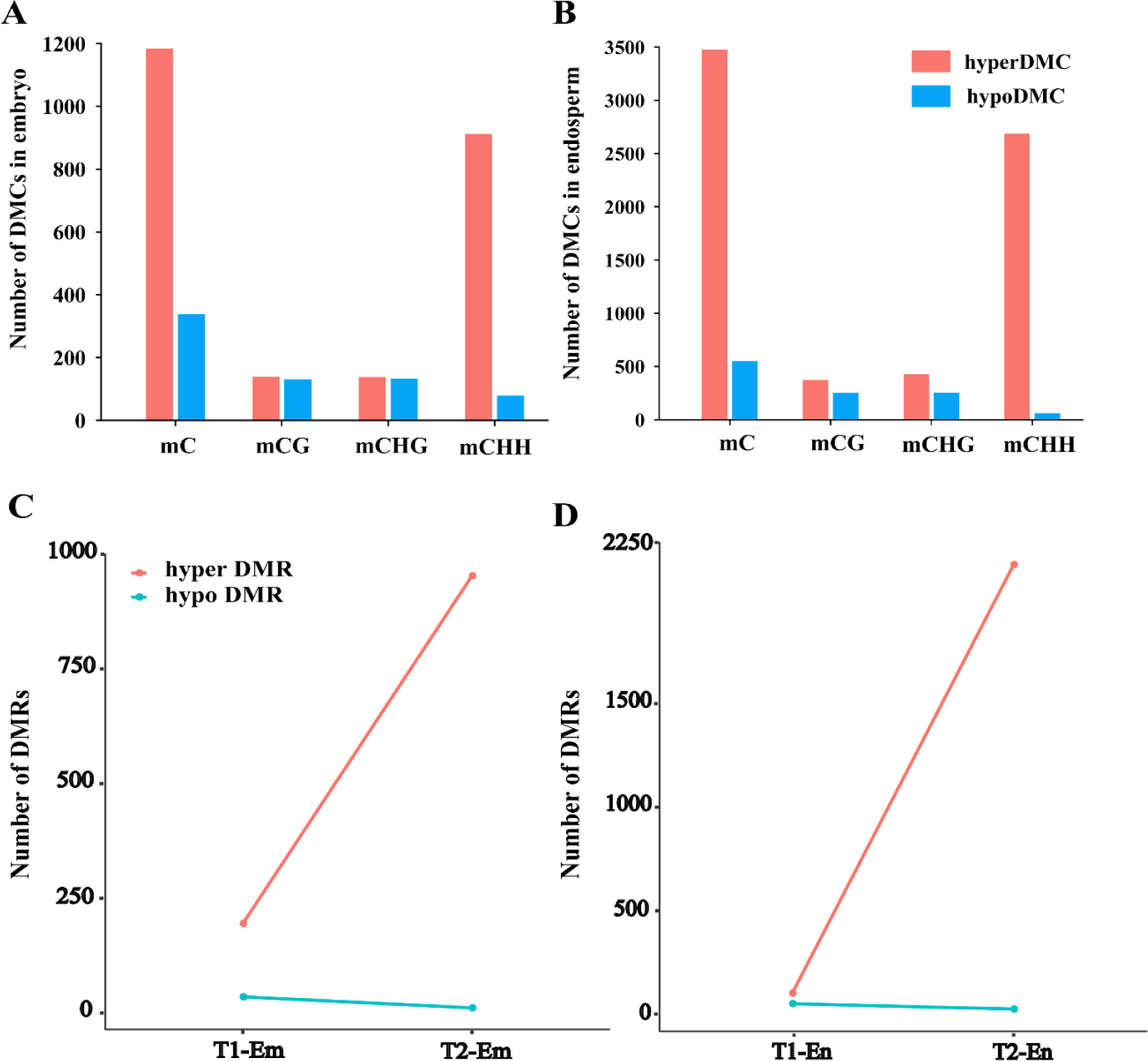
Numbers of the after-ripening process induced differentially methylated cytosines (DMCs) in the embryo (A) and endosperm (B), hyper- and hypo-differentially methylated regions (DMRs) in the embryo (C) and endosperm (D) in T2 relative to CK of *P. notoginseng* seeds.

To examine the difference between the dormant and dormancy-released seeds, we analyzed the differentially methylated cytosines (DMCs) of CK to T2. In the embryo, a total of 1517 DMCs were found in T2 compared to CK, and 65% of the hyper-DMCs occurred in the CHH sites (Fig. 4A). In the endosperm, a total of 4014 DMCs was identified in T2 relative to CK, among which 3469 were hyper-methylated (hyper-DMCs) and 545 were hypomethylated (hypo-DMCs). Furthermore, the hyper-DMCs in the mCG, mCHG, and mCHH were 611, 669, and 2734, respectively (Fig. 4B). It seemed that DMCs of *P. notoginseng* seeds showed a gradual increase during the after-ripening process. The CHH context is mainly responsible for the variations in DNA methylation, which makes up more than 98% of hyper-DMCs in the endosperm compared to less than 70% in the embryo. Therefore, DNA methylation in the genic regions of *P. notoginseng* seeds has a tissue-specific regulation pattern. To reveal the characteristics of dynamic changes in DNA methylation, a 200-bp sliding window was adapted to identify differentially methylated regions (DMRs). It showed that the hyper-DMRs identified in T1 and T2 relative to CK gradually increased (Fig. 4C, D). In contrast, there was no significant difference in the numbers of hypo-methylated regions (hypo-DMRs). In the embryo, we observed that the numbers of hyper-DMRs in T1 (195) and T2 (953) increased significantly compared to CK. Besides, it increased from 90 in T1 to 2,160 in T2 compared to CK in the endosperm (Fig. 4D). These results show an increase in global DNA methylation levels from CK to T2. To investigate whether the methylation variation occurs preferentially in any sequence context, DNA methylation levels of hyper-DMRs (Fig. 5) were examined in CG, CHG, and CHH, respectively. Hyper-DMRs displayed the same pattern of change, with an increase of DNA methylation from CK to T2 in the embryo and endosperm (Fig. S5). Especially, the CG methylation level shows a significant enhancement, and the hyper-methylation mainly occurs in CHH contexts. It suggested that the CHH hyper-methylation is not accompanied by CG demethylation in *P. notoginseng* seeds (Fig. 5). Overall, these results indicated that *P. notoginseng* seeds exhibit extensive DNA hyper-methylation during the after-ripening process.

**Figure 5.**
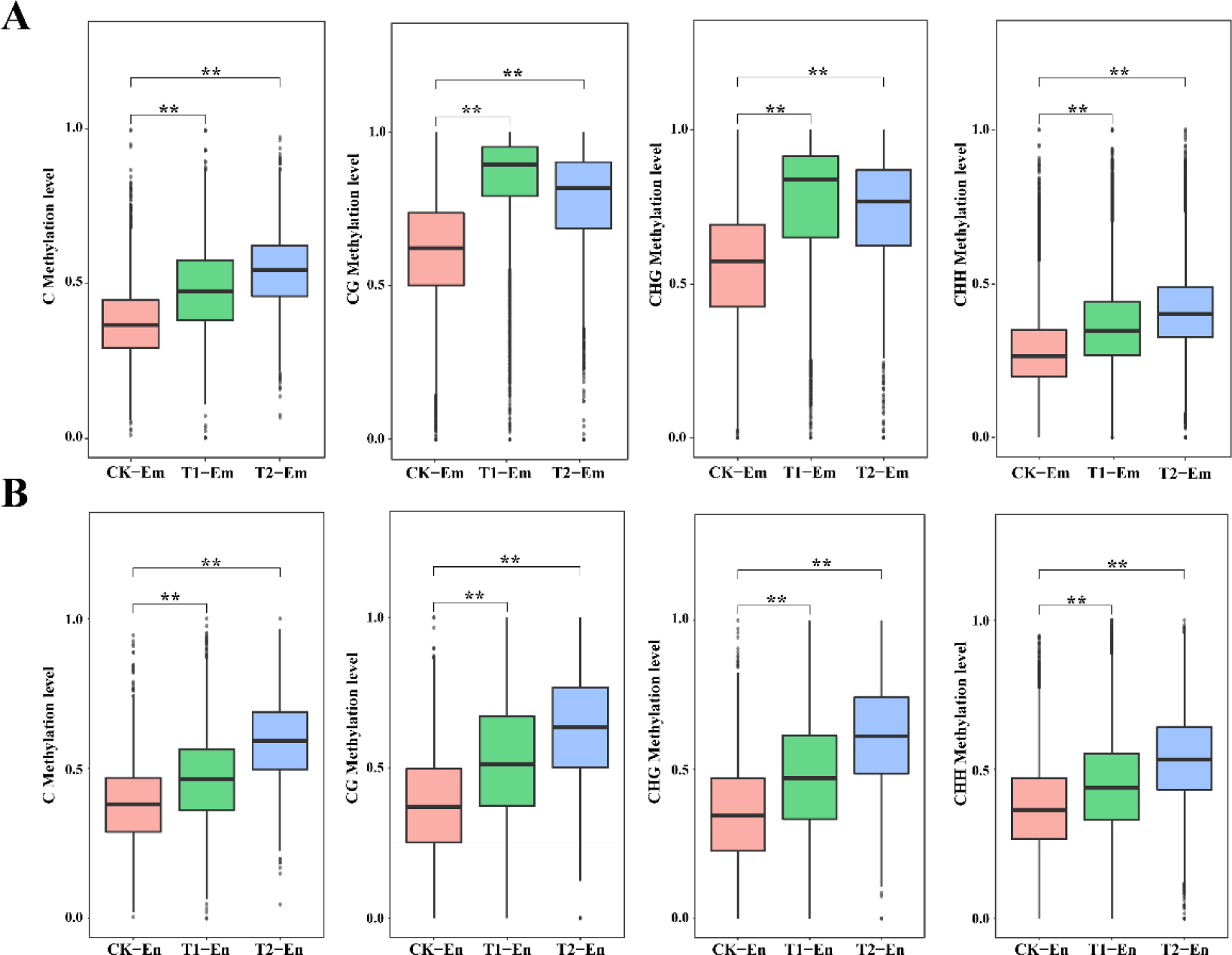
DNA methylation levels of hyper-DMRs in the embryo (A) and endosperm (B) during the after-ripening process. *adjusted *P* value < 0.05, **adjusted *P* value < 0.01.

### Increased DNA methyltransferase gene expression correlates with tissue-specific methylation elevation during recalcitrant *P. notoginseng* seed after-ripening process

In plants, RNA-directed DNA methylation mediates the establishment of DNA methylation and involves a number of proteins, small interfering RNAs (siRNAs), and scaffold RNAs. To elucidate the relationship between RdDM activity and DNA methylation, the expression level of the key genes in RdDM (*RDR2*, *DCL3*, *DCL4*, and *AGO4*) was examined (Fig. S6D). In general, most genes in RdDM exhibited a low transcription and there were no significant changes during the after-ripening process of *P. notoginseng* seeds, compared to CK. Among other higher transcription genes, most of them were up-regulated in T1, and they were subsequently down-regulated in the embryo. Similarly, the expression level in the endosperm was found to increase in T1 compared to CK (Fig. S6D). To further assess siRNA expression during the after-ripening process of *P. notoginseng* seeds, we analyzed the levels of 24-nt siRNA in the RdDM pathway. First, the 24-nt siRNAs were extracted from the miRNA data (Table S3), and we calculated the 24-nt siRNAs in the hyper-DMRs and hypo-DMRs from CK to T2. In the embryo, the levels of siRNA in the hyper-DMRs covered regions were significantly reduced (*P* < 0.05) during the after-ripening process. Similarly, the level of the 24-nt siRNAs was decreased in the hyper-DMRs of the endosperm from the CK to T2 (Fig. S6E). These results suggest a poor link between the enhanced DNA methylation and the changes in the RdDM activity during the after-ripening process.

DNA methylation and DNA demethylation work antagonistically to regulate the level of plant methylome. To reveal the mechanism of DNA hyper-methylation during the after-ripening process of *P. notoginseng* seeds, we also tested the expression of DNA methyltransferase and demethylase genes (Fig. S7, Additional file 2: Table S6). The genome-wide transcript profiles were generated in the embryo and endosperm of *P. notoginseng* seeds at the CK-T2 stages. It showed good consistency via PCA in three biological replicates at each stage (Fig. S8A). The *P. notoginseng* genome harbors sixteen methyltransferase homologs and seven demethylase homologs (Fig. S6A, B) using blastp, including eight *PnMET1*, three *PnCMT2*, one *PnCMT3*, four *PnDRM2*, four *PnROS1* and three *PnDME1* (Fig. 6, Fig. S6). For DNA demethylase genes, the four *AtROS1* orthologs were identified in *P. notoginseng* seeds (Fig. S6B), including Pno02G004385, Pno02G004391, Pno05G003816, and Pno09G001314. Moreover, we also identified three *AtDME1* orthologs, including Pno03G013947, Pno06G004027, and Pno07G001961. Of the seven *P. notoginseng* DNA demethylase genes, three *PnROS1* was significantly up-regulated in the embryo, whereas it was barely expressed in the endosperm. Especially, the *PnDME1* (Pno06G004027) in the embryo, and the *PnDME1* (Pno07G001961) in the endosperm were significantly down-regulated during the after-ripening process (Fig. 6B). The reduced expression of *PnDME1* in seeds is correlated with the gradually enhanced level of DNA methylation.

**Figure 6.**
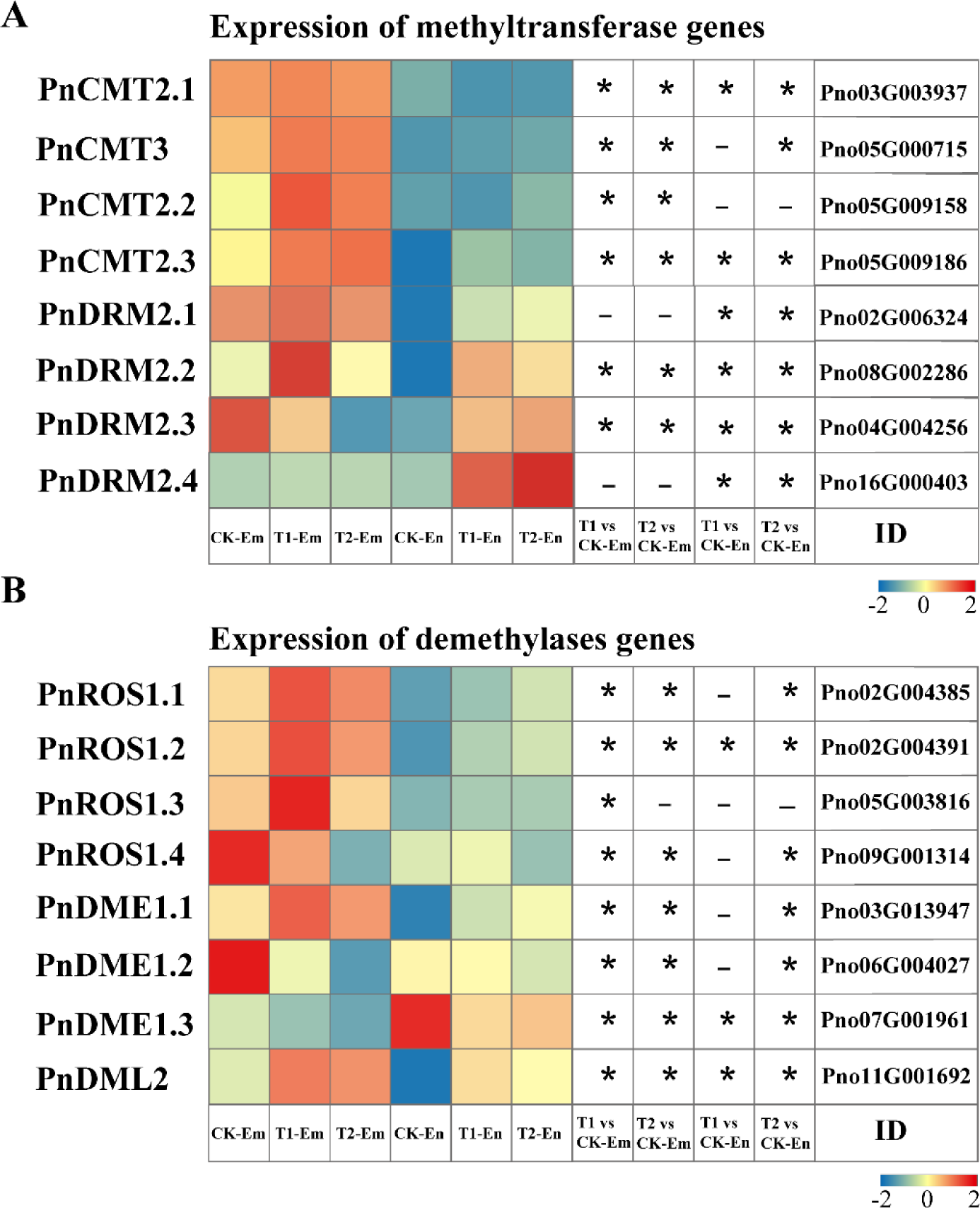
Expression of DNA methyltransferase and DNA demethylase genes in embryo and endosperm during the after-ripening process. Transcript levels of DNA methyltransferase (A) and DNA demethylase genes (B) during the after-ripening process. *adjusted *P* value < 0.05, as determined using the DESeq.

For DNA methyltransferase, in the seven *PnMET1* genes, four (Pno01G012352, Pno11G004052, Pno07G002989, Pno07G002988) were hardly expressed (FPKM < 2) in the endosperm. Among the other three genes, only one was up-regulated during the after-ripening process (Fig. S6C). In the embryo, it showed the same pattern as the endosperm that four genes show a low transcription. Especially, the DNA methyltransferase gene *PnCMT2* (Pno02G006324, Pno05G009158, Pno05G009186) and *PnCMT3* (Pno05G000715) in the embryo show an increased transcript levels during the after-ripening process, as the methyltransferase gene *PnDRM2* (Pno02G006324, Pno08G002286, Pno04G004256, Pno16G000403) in the endosperm did (Fig. 6). In contrast, the expression level of *PnCMT3* in the endosperm and *PnDRM2* in the embryo were down-regulated during the after-ripening process. Differential expression of the two DNA methyltransferases was observed between the embryo and endosperm tissues. Collectively, considering the expression changes of DNA methyltransferase and DNA demethylase genes, it supports that the DNA hyper-methylation of *P. notoginseng* seeds is most likely contributed by the enhanced expression of DNA methyltransferase genes.

To test the function of DNA methyltransferase, the TRV-mediated gene silencing was used to down-regulate *PnCMT2* in *P. notoginseng* seeds. It observed early germination in *PnCMT2* seeds compared to control seeds (Fig. 7). Moreover, compared to control seeds, the seed germination rate in *PnCMT2* was increased by 72.7% and 29.1% at 50 DAR and 60 DAR, respectively. These results support that the decreased expression of DNA methyltransferase *CMT2* leads to DNA hypo-methylation, thus contributing to recalcitrant seed development and after-ripening.

**Figure 7.**
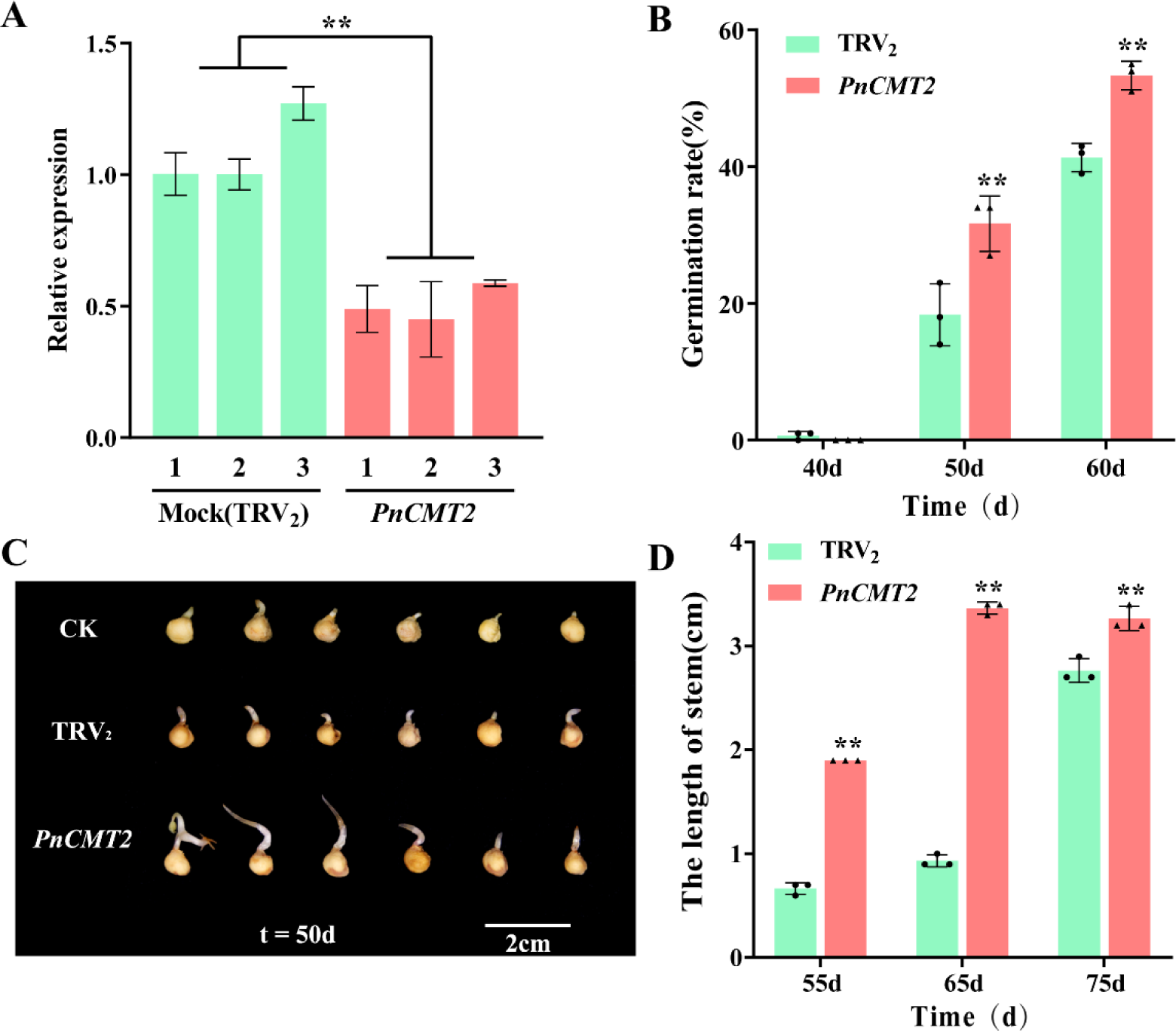
TRV_2_-induced gene silencing of *PnCMT2* in *P. notoginseng* seeds. (A)RT-qPCR analysis of *PnCMT2* in TRV_2_ control and TRV_2_: *PnCMT2* seeds. Changes in germination rate (B) and the stem length (D) of TRV_2_ control and TRV_2_: *PnCMT2* seeds during the after-ripening process. (C) Pictures of TRV_2_ control and TRV_2_: *PnCMT2* seeds after germination (t=50d). The values presented are the means ± SE (n=3). *adjusted *P* value < 0.05, **adjusted *P* value < 0.01.

### DNA methylation increase is linked to gene expression in *P. notoginseng* seeds

To further investigate the effects of DNA methylation on gene expression of *P. notoginseng* seeds, we compared the transcriptomes for embryo and endosperm in CK to T2. It showed a high consistency among the biological replicates (Fig. S8A). For each sample, 86.54% ∼ 89.18% of the reads in the 18 libraries were mapped by alignment with the reference genome (Table S2). The distribution of the log2 (FPKM+1) had a relatively high gene expression (Fig. S8B). Furthermore, we identified 6494, 9532, 9377, and 10270 differentially expressed genes (DEGs, adjusted *P* value < 0.01) in T1-Em, T2-Em, T1-En, and T2-En relative to CK, respectively (Fig. 8A). We found that DEGs identified in the different stages had a high overlap in Venn diagrams (Fig. 8B). The number of DEGs in the endosperm was significantly greater than that of DEGs in the embryo. Moreover, the up DEGs and down DEGs of seeds were gradually increased from T1 to T2, suggesting that overall gene expression levels are gradually changed during the recalcitrant seed after-ripening process.

**Figure 8.**
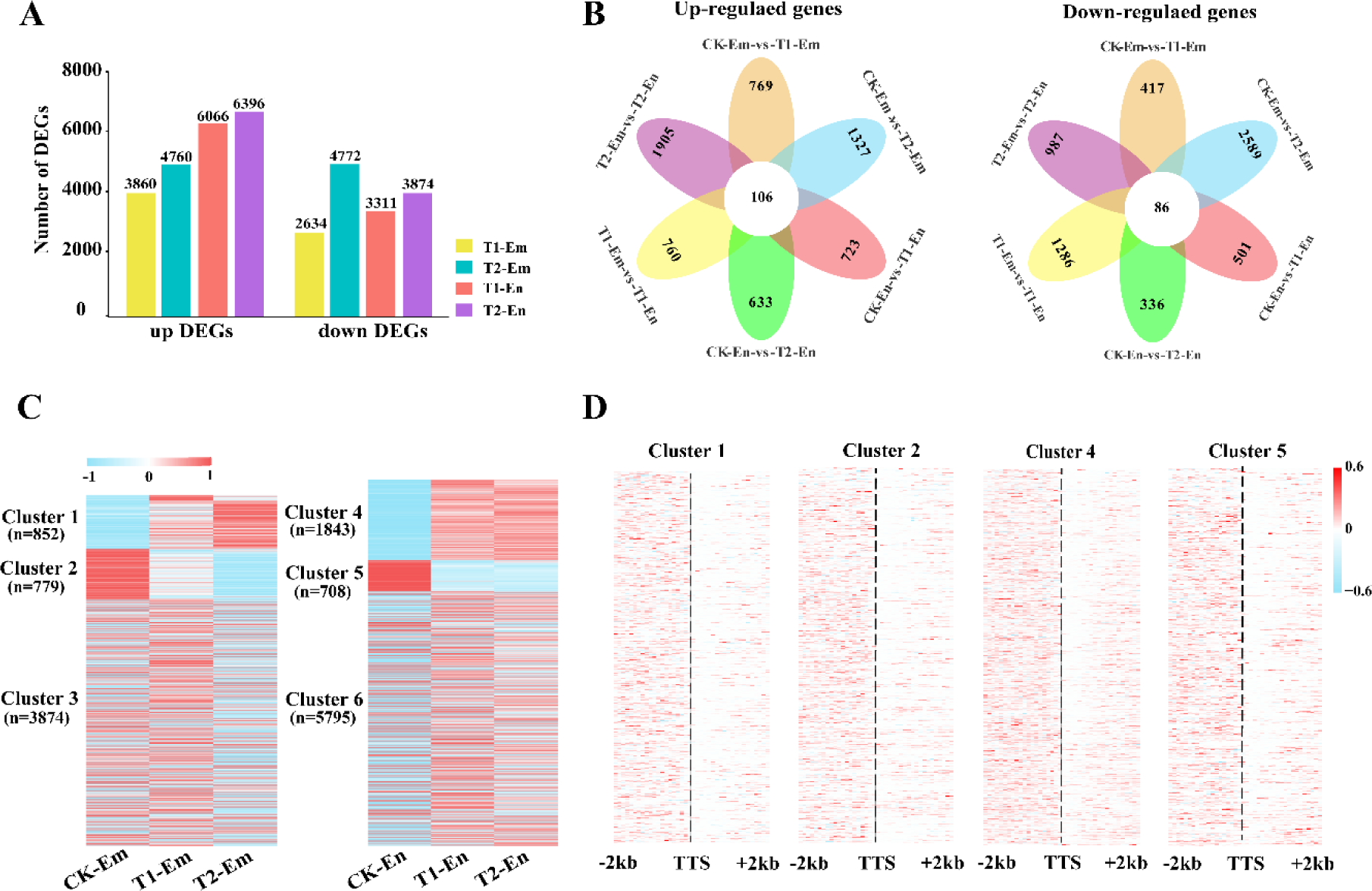
Characterization of the expression and DNA methylation of hyper-DMR-associated genes. (A) Histogram showing the numbers of DEGs in T1 and T2 relative to CK in embryo and endosperm. (B) Venn diagrams show overlaps among the up-regulated and down-regulated DEGs. (C) Heatmaps showing the expression levels of the six clusters of hyper-DMR-associated genes during the after-ripening process. (D) DNA methylation changes of the four clusters of genes in T2 vs CK in the embryo and endosperm. The sequences flanking genes were aligned by transcription start sites (TTS), and DNA methylation levels for each 100-bp interval were plotted. The dashed line marks transcription start sites.

To reveal the relationship between DNA methylation changes and gene dynamic regulation during the after-ripening process, we examined the hyper-DMRs-associated genes because *P. notoginseng* seeds undergo overall DNA hyper-methylation. It found a very heavy bias for hyper-DMRs containing methylation changes in the CHH context (Fig. 4C). Compared to CK, we observed a total of 953 and 2160 hyper-DMRs in T1 or T2 of the embryo and endosperm (Additional file 1: Table S5). Among these hyper-DMRs, we found 5505 and 8436 hyper-DMRs-associated genes in the embryo and endosperm, respectively, and they were differentially expressed in at least one of the stages (Fig. 8B). Next, we found 852 up-regulated DEGs (cluster 1), 779 down-regulated DEGs (cluster 2), and 3874 non-significantly changed DEGs in the embryo (cluster 3). Similar to the embryo, there are 1843 up-regulated DEGs (cluster 4), 708 down-regulated DEGs (cluster 5), and 5795 non-significantly changed DEGs (cluster 6) in the endosperm (Additional file 3: Table S7). The expression of cluster 1 and cluster 4 was gradually increased during the after-ripening process, while the expression of cluster 2 and cluster 5 was decreased (Fig. 8C). It may be shown that cluster 3 and cluster 6 genes show a small change due to barely changes in DNA methylation compared to cluster 1, cluster 2, cluster 4, and cluster 6.

To understand the importance of DNA methylation in regulating gene expression, we analyzed the DNA methylation changes of up-regulated DEGs, down-regulated DEGs (cluster 5), and non-significant changes in DEGs in the embryo and endosperm. It observed that the DNA methylation of the up- and down-regulated DEGs undergoes a global increase from CK to T2, especially in promoter regions (Fig. 8D). Furthermore, we examined the promoter methylation levels of the six clusters of genes during the after-ripening process (Fig. S8C). It found that the DNA methylation of clusters 1 and 2 in the embryo, and clusters 4 and 5 in the endosperm increased gradually from CK to T2. Similarly, the expression level of these genes also changed gradually during the after-ripening process (Fig. 8C). Thus, we further analyzed the association between DNA methylation and gene expression in the embryo and endosperm. As shown in Fig. S9, we calculated the correlation coefficient between the expression level and the promoter DNA methylation from CK to T2. It showed that the gene expression level and the DNA methylation level show a strong positive correlation (R > 0.85) in cluster 1 and cluster 4 (Fig. S9A). In contrast, the gene-expression level and the DNA methylation of the cluster 2 genes and cluster 5 genes showed a highly negative relationship (R < -0.85). These results support that DNA methylation may play a positive role in regulating the expression of many genes during the recalcitrant seed after-ripening process.

### Genes associated with DNA hyper-methylation

To assess how hyper-DMRs-associated gene regulates recalcitrant seed after-ripening and development, the Gene Ontology (GO) analysis and KEGG analysis for hyper-methylated associated DEGs were performed in the embryo and endosperm, respectively. For the up-regulated DEGs in the embryo (cluster 1), the analysis found that 29 genes involved in cluster 1 are enriched in the hormone-mediated signaling pathway, 29 genes are enriched in plant organ development, and 9 genes are annotated in the ethylene-activated signaling pathway (Additional file 4: Table S8). Plant hormone is important for seed development and after-ripening, which can be regulated by embryo maturation programs during the after-ripening process. Consistently, our KEGG analysis found that hyper-DMRs-associated genes (cluster 1 and cluster 2) were enriched in the plant hormone signal transduction, and MAPK signaling pathway-plant (Fig. S10A), suggesting that after-ripening-induced DNA hyper-methylation regulates the expression of plant hormone signaling-related genes (Fig. 9). It is known that DNA methylation, especially at promoters, is normally considered to lead transcriptional silencing. For the down-regulated DEGs in the embryo (cluster 2), genes involved in embryo development, embryonic axis specification, gene silencing, and seed development are enriched (Additional file4: table S8). It is in agreement with the potential role of DNA methylation in gene silencing. As shown in Fig. 9B, we identified 187 genes that are annotated in the anatomical structure development, 60 genes are annotated to the fatty acid metabolic process, and 17 genes are enriched in developmental maturation (cluster 4). For the down-regulated DEGs in the endosperm, 103 responses to stress-related genes, 13 regulation of post-embryonic development-related genes, and 7 seedling development-related genes were enriched in cluster 5. Moreover, the KEGG analysis showed that hyper-DMRs-associated genes (cluster 4 and cluster 5) were enriched in fatty acid degradation and pyruvate metabolism (Fig. S10A). According to our analysis, it appears that DNA hyper-methylation contributes to the after-ripening of *P. notoginseng* seeds.

**Figure 9.**
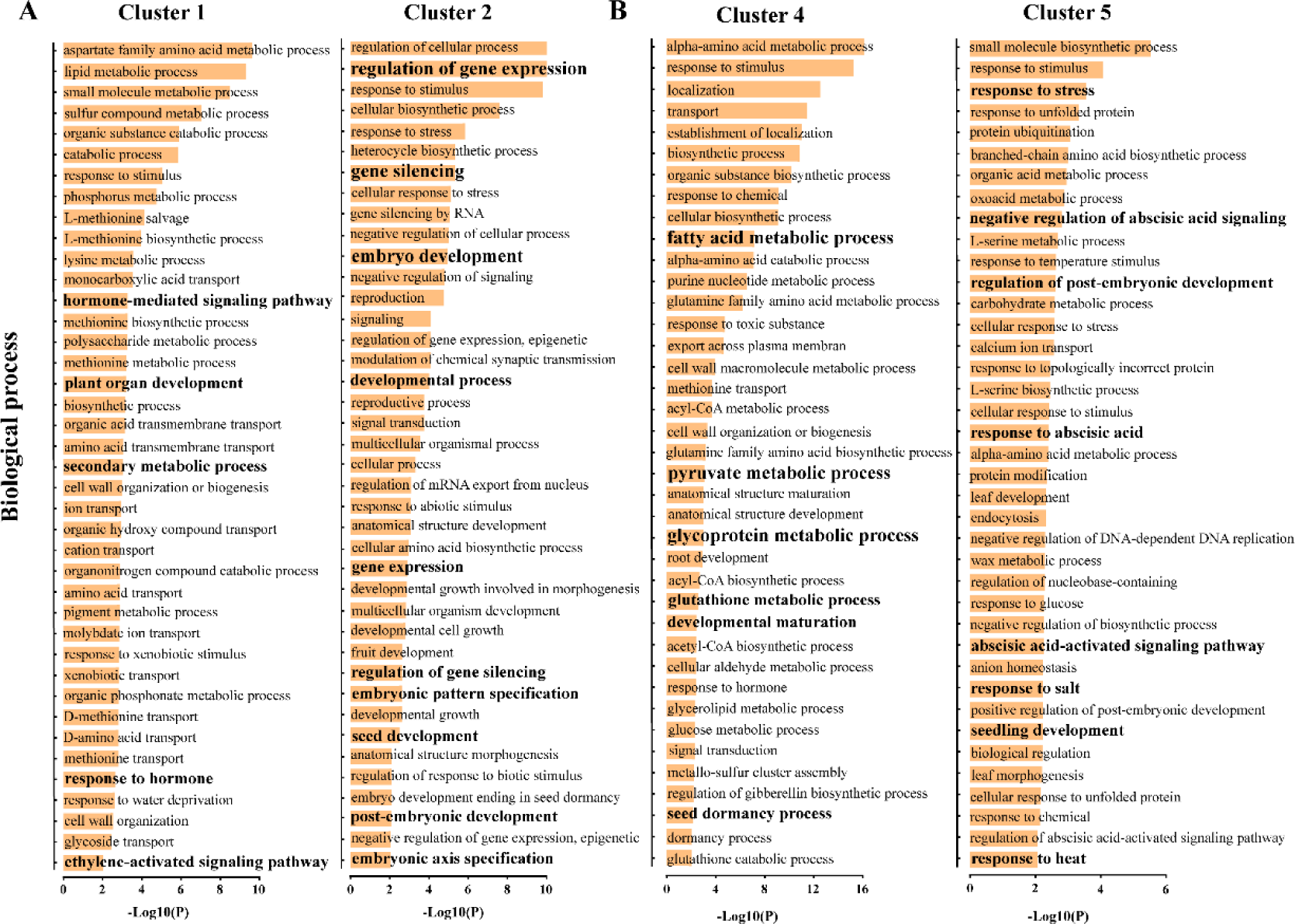
Functions of hyper-DMR-associated DEGs. GO analysis of hyper-methylated DEGs in the embryo (A) and endosperm (B). GO terms enriched in hyper-methylated DEGs were illustrated.

To examine the importance of DNA methylation for *P. notoginseng* seeds during the after-ripening process, the DNA methylation inhibitor 5-azacytidine and DNA methylation promoter hydroxyurea were applied to *P. notoginseng* seeds at 0 DAR. As shown in Fig. 10A, the 5-azacytidine-treated seeds germinated prematurely compared to the mock. In contrast, the germination of the hydroxyurea-treated seeds was inhibited slightly. To analyze whether the inhibitor and promoter change DNA methylation level, the DNA methylation of two up-DEGs (Pno05G014320, Pno08G004112) and two down-DEGs (Pno08G004112, Pno08G004112) in mock and treated seeds was measured (Fig. 10B). It showed that the promoter regions of the four genes are hyper-methylated in T2 compared to CK in mock seeds (Fig. S10), this is agreed with the observation that the DNA methylation globally increased during the after-ripening process of *P. notoginseng* seeds. In the 5-azacytidine-treated seeds, the McrBC-qPCR assay found that the promoter regions of four genes were decreased in the DNA methylation levels. In contrast, it found that DNA methylation increased in the hydroxyurea-treated seeds compared to the mock at 50 DAR. To further examine the effect of 5-azacytidine and hydroxyurea on the gene expression levels, we assessed the expression of the four genes. Compared to the mock, the up-DEGs in the embryo and endosperm were down-regulated in the 5-azacytidine-treated samples (Fig. 10C), whereas they were up-regulated in the hydroxyurea-treated samples. For the down-DEGs, the DNA hypo-methylation caused a rise in expression level in the 5-azacytidine-treated seeds, supporting the fact that DNA hyper-methylation represses the expression of two genes. Overall, after-ripening-mediated DNA hyper-methylation has a critical role in regulating the expression of recalcitrant seed development-related and after-ripening-related genes.

**Figure 10.**
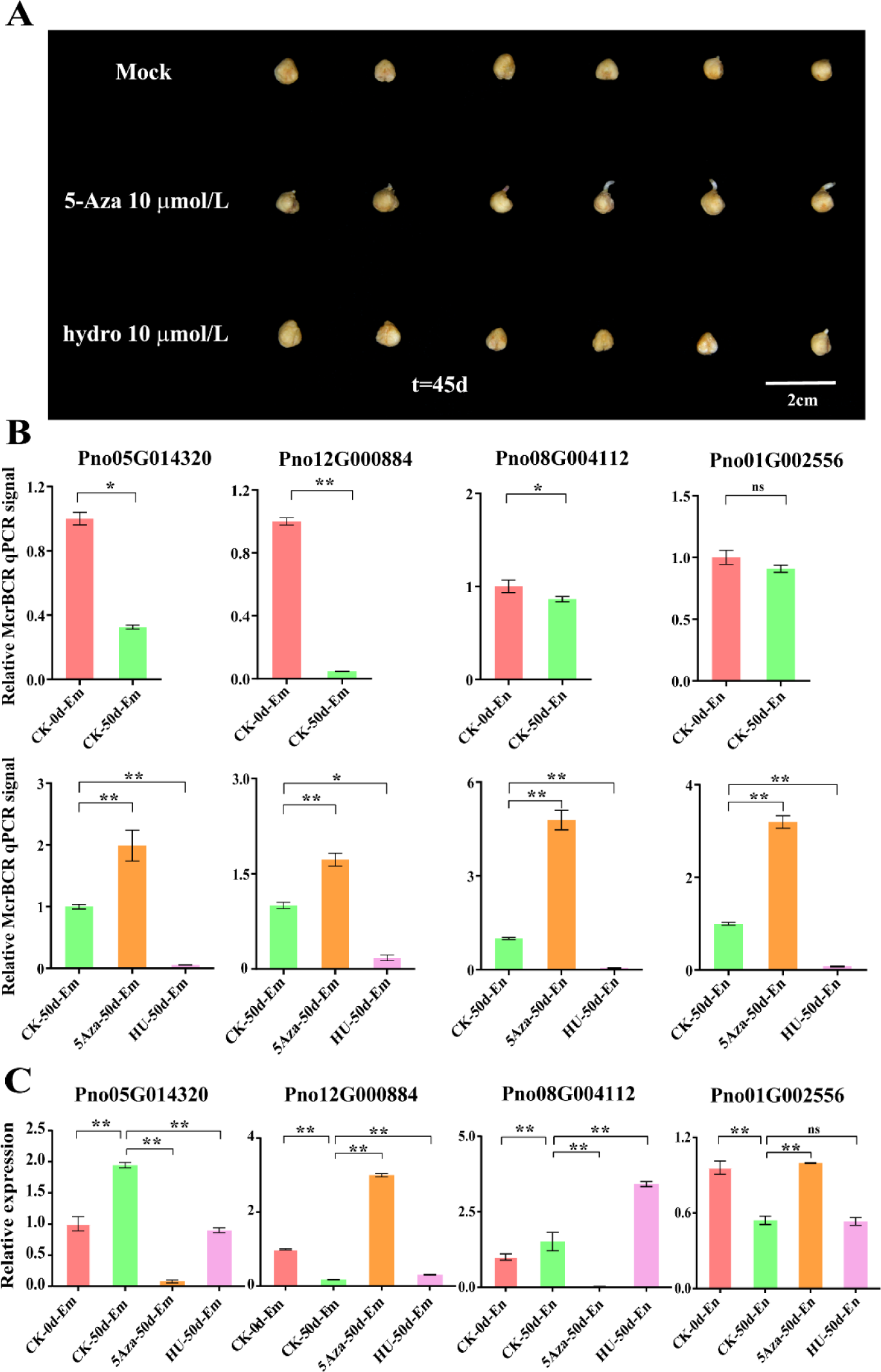
The significance of DNA methylation for *P. notoginseng* seeds during the after-ripening process. (A) Pictures of *P. notoginseng* seeds that were treated with 5-azacytidine (5-Aza), DNA methylation promoter hydroxyurea (HU), and mock (ddH_2_O). (B) Gene expression of Pno05G014320, Pno12G000884, Pno08G004112, and Pno01G002556 in untreated CK and T2 seeds, and in CK, 5-Aza, and HU-treated seeds. (C) DNA methylation levels of promoter regions of Pno05G014320, Pno12G000884, Pno08G004112, and Pno01G002556. McrBC-qPCR analysis was performed in CK, T2, 5-Aza, and HU-treated samples. Lower qPCR signals indicate higher methylation levels. \**P* < 0.05; \*\**P* < 0.01. Error bars indicate means ± SD.

## Discussion

### It is hyper-methylated in the endosperm genome of *P. notoginseng* seeds

DNA methylation is an essential epigenetic modification in plant genomics(Zhang *et al*., 2018). The characterization of DNA methylation is valuable for investigating the regulatory relationship between genomes and plant phenotypes(Xiao *et al*., 2020; Tian *et al*., 2021). Whole-genome bisulfite sequencing with a single base sequencing depth of 20-fold has confirmed that average methylation levels of mCG, mCHG, and mCHH are 24%, 6.7%, and 1.7% in *Arabidopsis*, respectively(Cokus *et al*., 2008). It has also been recorded that H3K27me3 and DNA methylation exist in transcriptionally inactive genes and repetitive elements in maize seeds(Wang *et al*., 2009). To characterize the DNA methylomes of *P. notoginseng* seeds, genome-wide maps at a single-base resolution in DNA methylation were performed (Fig. S1). The average coverage of our sequenced methylomes was ∼30-fold. The depth of these sequenced methylomes is consistent with those of the sequenced methylomes in *Arabidopsis*(Lister *et al*., 2008) and maize(Zhang *et al*., 2014), suggesting that the quality of DNA methylomes obtained from bisulfite sequencing shows a sufficient level for further analysis.

The significance of DNA methylation is mainly reflected in the strict control of DNA methylation in different tissues or cells(Slotkin *et al*., 2009; Ibarra *et al*., 2012). An analysis of DNA methylation has revealed that average methylation levels of CG, CHG, and CHH are 20.9%, 8.9%, and 2.8% in the endosperm of *Arabidopsis* seeds, being lower than those in its embryo (26.9% CG, 10.6% CHG, and 4.4% CHH)(Hsieh *et al*., 2009). Likewise, the DNA methylation level in rice endosperm is less than those in the embryo, and the endosperm hypo-methylation activates the expression of starch-synthesizing enzymes and genes coding for storage proteins in rice endosperm(Zemach *et al*., 2010). In *P. notoginseng* seeds (CK), we found that the methylation level is 82% for CG, 75% for CHG, and 15% for CHH in the embryo, and 90% for CG, 82% for CHG and 22% for CHH in the endosperm (Fig. S4), indicating that the methylation levels of three sequence contexts in the embryo and endosperm of *P. notoginseng* seeds are significantly higher than those in the embryo and endosperm of *Arabidopsis* seeds(Chen, M *et al*., 2018) and rice seeds(Rodrigues *et al*., 2013). Interestingly, DNA methylation levels in different sequence contexts are stable across the species. CG context showed the highest level of methylation while the CHH context did the lowest level (Fig. S4). This indicates that a relatively consistent DNA methylation exists throughout the evolution of different species. Moreover, the embryo genome was hypo-methylated compared with the endosperm genome in our data, and this is different from those observed in the PD-typed seeds of maize(Lu *et al*., 2015), rice(Zemach *et al*., 2010), and castor (*Ricinus communis*)(Xu *et al*., 2016). Thus, we suppose that a relative hypo-methylation might be more beneficial for the incompletely developed embryo to accomplish the after-ripening in the MPD-typed seeds.

### DNA methylation is dramatically enhanced by the upregulation of DNA methyltransferase during the process of recalcitrant seed after-ripening

Besides hormones and transcription factors, DNA methylation is considered to be one of the essential factors that influences fruit ripening and seed development(Zhong *et al*., 2013; Liu *et al*., 2015; Chen, M *et al*., 2018). CHH methylation programmatically increases from the glob stage through the dormancy in the PD-typed soybean and *Arabidopsis* seeds, but drops precipitously within the germinating seedling(Lin *et al*., 2017). Likewise, a global increase was observed in CHH methylation (Fig. 3), especially in the embryo during the after-ripening process (from 31.8 to 34.8%), this is also observed during the orange fruit development and ripening(Huang *et al*., 2019). These results indicate that DNA hyper-methylation also occurs in the MPD-typed seeds with recalcitrant characteristics. In tomatoes, a global CHH hyper-methylation is accompanied by CG and CHG hypo-methylation during the process of ripening(Zhong *et al*., 2013). Similarly, our analysis found that the enhancement in DNA methylation is mostly due to the increase in CHH methylation (Fig. 4), whereas CHH hyper-methylation is not accompanied by CG and CHG hypo-methylation during the after-ripening of *P. notoginseng* seeds (Fig. S4). These results agree with the fact that in the soybean and *Arabidopsis*, there are no significant global changes in methylation of CG and CHG context during the seed developmental period. Overall, CHH hyper-methylation contributes to the global increase of DNA methylation during the process of recalcitrant seed after ripening.

DNA demethylases and DNA methyltransferases could dynamically regulate DNA methylation status. In *Arabidopsis*, DNA demethylation is mediated by ROS1, which removes cytosine bases by cutting DNA strands and completes the demethylation process(Deleris *et al*., 2016). During the strawberry fruit ripening, DNA methylation is dramatically reduced by the down-regulation of RdDM, and the DNA methylation inhibitor causes an early ripening phenotype(Cheng *et al*., 2018). However, we observed that only a few genes in the RdDM pathway are up-regulated during the after-ripening process of *P. notoginseng* seed (Fig. S6D). Meanwhile, it showed that the expression level of three *PnROS1* significantly increased in the embryo (Fig. 6). And the ripening-induced hyper-DMRs in *P. notoginseng* seeds were enriched with 24-nt siRNAs (Fig. S6E). In *Arabidopsis*, *AtROS1* interferes with RdDM to avoid DNA hyper-methylation, and 24-nt small interfering RNAs (siRNAs) that cause RdDM are enriched in regions targeted by *AtROS1*(Zhang *et al*., 2022). The expression pattern between *PnROS1* and RdDM activity suggests that the increased expression of *PnROS1* could antagonize RdDM to arrest DNA hyper-methylation during the *P. notoginseng* seed after-ripening process. Thus, we conclude that the increased DNA methylation may be independent of RdDM activity and *PnROS1*-mediated DNA demethylation. During the orange fruit ripening, the reduced expression of *CsDME* and *CsDML* leads to DNA hyper-methylation(Tang *et al*., 2016). We then determined whether the expression level of *DME* contributes to the hyper-methylation of the embryo and endosperm in *P. notoginseng* seeds. After the comparison of CK with T2, we found that only one *PnDME* is significantly up-regulated during the after-ripening process (Fig. 6B). Especially, all of the *PnCMT2s* in the embryo and *PnDRM2s* in the endosperm significantly increased during the after-ripening process (*P*<0.05), and this is also similar to the trend of the after-ripening-induced hyper-methylation (Fig. 6A). Consistently, TRV-mediated *PnCMT2* silencing seeds (Fig. 7) have shown that the elevated DNA methylation might be regulated by *PnCMT2* factors. Our results partly support the findings that the increased DNA methylation at the CHH context of the *Arabidopsis* embryo is dependent on a sustained increase in *AtCMT2*(Lin *et al*., 2017). Taken together, our study reveals that the enhanced DNA methylation is most likely caused by the increased expression of DNA methyltransferase during the after-ripening process of recalcitrant seeds.

### DNA hyper-methylation regulates the after-ripening-related genes in seeds

In plants, DNA methylation might happen at the promoter or within the transcribed gene body, and DNA methylation in the promoter usually represses gene transcription(He *et al*., 2021; Wang *et al*., 2022). The ripening-induced hyper-methylation in the orange fruit represses the expression of cell wall organization-related and photosynthesis-related genes. In *Arabidopsis* seeds, hyper-methylation enhances chromatin packaging to prevent unfavorable gene expression during seed dormancy(Bouyer *et al*., 2017). During the after-ripening process of *P. notoginseng* seeds, we found that DNA hyper-methylation enhancement occurs in the promoter regions of genes in the embryo and endosperm (Fig. 8). Consistently, it was discovered that a strong correlation exists between the expression level of hyper-DMRs-associated genes and the DNA methylation level (Fig. S9), suggesting that DNA hyper-methylation regulates the expression of hyper-DMRs genes during the after-ripening process. Furthermore, hyper-methylation in the embryo caused a reduced expression of genes responsible for embryo development, embryonic pattern specification, and seed development (Fig 9). These findings are in agreement with the silencing role of DNA methylation when occurs at the transcriptional level in rice and *Arabidopsis*(Stroud *et al*., 2013; Wang *et al*., 2022). On the other hand, the down-regulated expression genes in the endosperm were enriched in response to heat and stress, negative regulation of abscisic acid signaling, and seedling development (Fig. 9). Abscisic acid (ABA) inhibits or delays seed germination and embryo development(Yan & Chen, 2017; Quesada, 2021), and it is a positive regulator for seeds responding to dehydration(Smolikova *et al*., 2021). This suggests that the inhibition by DNA hyper-methylation might facilitate seed dormancy and make seeds sensitive to dehydration. Likewise, it showed that the DNA methylation inhibitor, 5-azacytidine, facilitates seed development and germination (Fig. 10), suggesting that DNA hyper-methylation may be critical for making seeds maintain a dormant status during the after-ripening process. Therefore, we propose that the repressed role by DNA hyper-methylation may contribute to sustaining seed dormancy when the embryo is incompletely developed during the after-ripening process.

DNA methylation, especially DNA methylation in gene promoters, represses gene expression. However, recent studies have suggested that DNA methylation also plays a positive role in gene expression. An MITE insertion in the promoter of the *HaWRKY6* in sunflower (*Helianthus annuus*) triggers DNA methylation, consequently improving the expression of the *HaWRKY6* in cotyledons(Gagliardi *et al*., 2019). It has been established that DNA methylation in the ROS1 promoter accelerates the expression of gene in *Arabidopsis*(Lei *et al*., 2015). Similarly, our results have shown that DNA hyper-methylation is also correlated with gene activation during the after-ripening process of *P. notoginseng* seeds (Fig. S9). In the embryo, hyper-DMRs-related genes (Cluster 1) were enriched in the hormone-mediated signaling pathway, plant organ development, and ethylene-activated signaling pathway (Fig. 9), and these pathways are closely related to seed development(Shu *et al*., 2016; Matilla, 2020). The activation of DNA hyper-methylation is consistent with the fact that RdDM positively regulates DWARF14 (D14) at a nearby MITE in rice(Wang *et al*., 2020). Expectedly, the endosperm provides nourishment to the embryo during early embryogenesis, and it degenerates and remains as a vestigial cell layer in the mature seed(Palovaara *et al*., 2013). In this study, we found that DNA hyper-methylation in the endosperm of *P. notoginseng* seed is significantly correlated with the increased expression of genes involved in the fatty acid metabolic process, pyruvate metabolic process, and glycoprotein metabolic process (Fig. 9). Similarly, the DNA hyper-methylation in orange ripening actives the expression of genes in ABA response and signaling pathway(Huang *et al*., 2019). Taken together, a critical role of DNA hyper-methylation in the activation of gene expression has been observed in the recalcitrant *P. notoginseng* seeds. Hyper-methylation might facilitate the activation of genes involved in the hormone-mediated signaling pathway during embryo development, and DNA hyper-methylation in the endosperm may activate energy metabolism-related genes to provide nourishment for the embryo development of *P. notoginseng* seeds.

## Conclusion

Our work firstly illustrates a genome-wide survey of DNA methylation and its association with gene expression during the after-ripening process of recalcitrant seeds. We propose a hypothetical model to explain how DNA methylation is changed during the after-ripening process of recalcitrant seed (Fig. 11). It suggests that CHH hyper-methylation regulates the after-ripening and dormancy of recalcitrant seeds. The high expression level of DNA methyltransferase such as *PnCMT2* in the embryo, and *PnDRM2* in the endosperm leads to DNA hyper-methylation, thus modifying the transcriptional status of genes associated with hyper-DMRs. Among them, transcriptional repression contributes to the seed dormancy and after-ripening. Meanwhile, DNA hyper-methylation activates gene expression in the recalcitrant seeds with after-ripening. This activation alters the hormone-mediated signaling pathway and energy metabolism, and thus would largely contribute to embryo development and after-ripening process. The regulation mechanisms highlight the essential role of DNA methylation in mediating the after-ripening process of recalcitrant seeds and provide us with a more integrative view of the epigenetic regulation of MPD-typed seed dormancy in plants.

**Figure 11.**
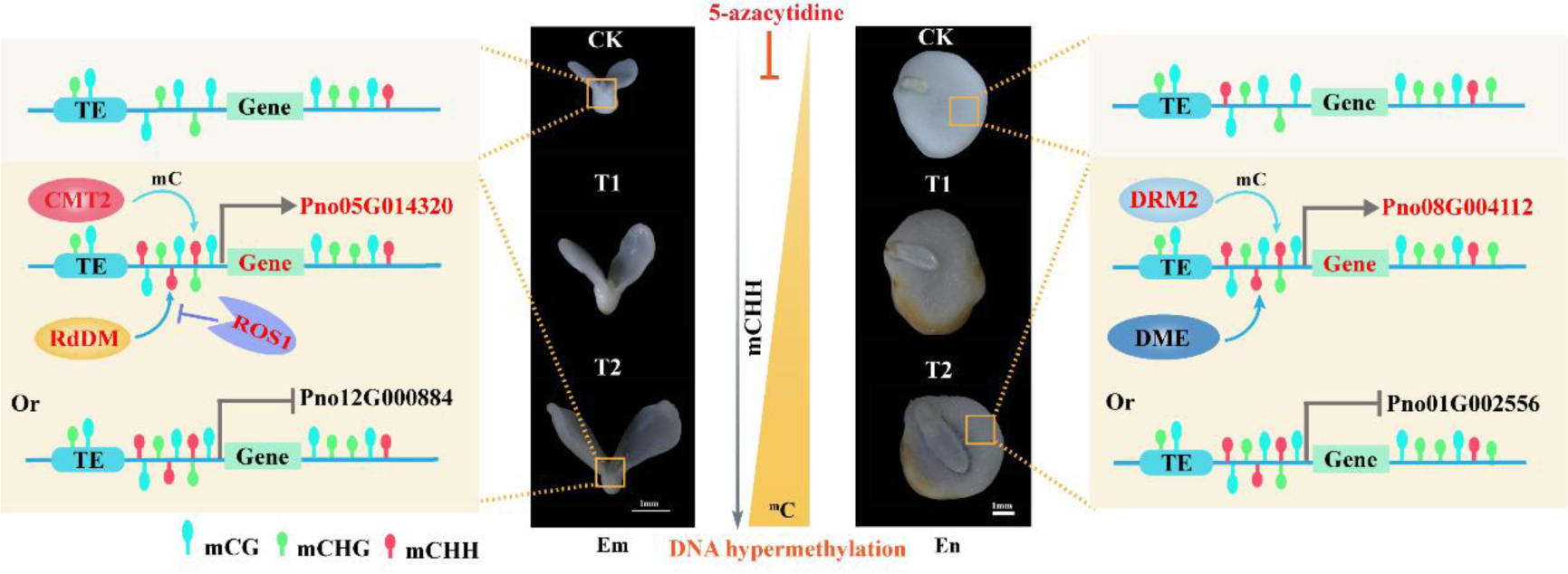
A hypothetical model to explain the DNA methylation pattern during the after-ripening process in the embryo and endosperm of *P. notoginseng*. Genes marked in red indicate that the genes were hyper-methylation-induced, and likewise, black ones indicate that the genes were hyper-methylation-repressed. Arrows and blunted lines designate positive and inhibitory interactions, respectively.

## Availability of data and materials

The raw sequencing data from this study have been deposited in the Genome Sequence Archive at BIG Data Center (https://bigd.big.ac.cn/), Beijing Institute of Genomics (BIG), Chinese Academy of Sciences, under the accession number: CRA012205, CRA012204, and CRA012206. Hoo & Tseng first undertook the formal identification of the plant material *Panax notoginseng* (Burkill) (Journal of Systematics and Evolution 11: 435, 1973) in Flora of China.

## Acknowledgments

This work was supported by the National Natural Science Foundation of China (32160248 and 81860676), the Major Special Science and Technology Project of Yunnan Province (202102AA310048), the National Key Research and Development Plan of China (2021YFD1601003), and the Innovative Research Team of Science and Technology in Yunnan Province (202105AE160016).

## Author contributions

J. C. directed the experiment and made suggestions for the writing of the manuscript. N. G. wrote the manuscript, and J. J., N. G. participated in most of the experiments. Y. W., C. L., and M. H. analyzed the relevant experimental data. All authors contributed to the article and approved the submitted version.

## Competing Interests

The authors declare no competing interests.

## Ethics approval and consent to participate

Not applicable.

## Authors’ information

^1^ College of Agronomy & Biotechnology, Yunnan Agricultural University, Kunming, Yunnan 650201, China. ^2^ The Key Laboratory of Medicinal Plant Biology of Yunnan Province, Yunnan Agricultural University, Kunming, Yunnan 650201, China. ^3^ National & Local Joint Engineering Research Center on Germplasm Innovation & Utilization of Chinese Medicinal Materials in Southwestern China, Yunnan Agricultural University, Kunming, Yunnan 650201, China.

